# Imeglimin Amplifies Glucose-Stimulated Insulin Release from Diabetic Islets via a Distinct Mechanism of Action

**DOI:** 10.1101/2020.10.20.346841

**Authors:** Sophie Hallakou-Bozec, Micheline Kergoat, Pascale Fouqueray, Sébastien Bolze, David E. Moller

## Abstract

Pancreatic islet β-cell dysfunction is characterized by defective glucose-stimulated insulin secretion (GSIS) and is a predominant component of the pathophysiology of diabetes. Imeglimin, a novel first-in-class small molecule tetrahydrotriazine drug candidate, improves glycemia and GSIS in preclinical models and clinical trials in patients with type 2 diabetes; however, the mechanism by which it restores β-cell function is unknown. Here, we show that Imeglimin acutely and directly amplifies GSIS in islets isolated from rodents with Type 2 diabetes via a mode of action that is distinct from other known therapeutic approaches. The underlying mechanism involves increases in the cellular nicotinamide adenine dinucleotide (NAD^+^) pool – potentially via the salvage pathway and induction of nicotinamide phosphoribosyltransferase (NAMPT) along with augmentation of glucose-induced ATP levels. Further, additional results suggest that NAD^+^ conversion to a second messenger, cyclic ADP ribose (cADPR), via cyclic ADP ribose hydrolase (CD38) is required for Imeglimin’s effects in islets, thus representing a potential link between increased NAD^+^ and enhanced glucose-induced Ca2+ mobilization which - in turn - is known to drive insulin granule exocytosis. Collectively, these findings implicate a novel mode of action for Imeglimin that explains its ability to effectively restore β-cell function and provides for a new approach to treat patients suffering from Type 2 diabetes.

## INTRODUCTION

Type 2 diabetes (T2DM) is characterized by insulin resistance plus β-cell dysfunction (1). Existing therapies may only be partially effective or not well tolerated (1). Glucagon-like peptide receptor (GLP1) agonists act on β-cells to amplify GSIS (2). However, these agents are peptides with limited oral bioavailablity and are usually administered parenterally. Therefore, the pursuit of newer therapies, in particular small molecules which could function to reverse β-cell dysfunction, is warranted.

Imeglimin is a novel oral antidiabetic drug to treat Type 2 diabetes. Its novel structure and proposed mechanism of action establishes the first in a new tetrahydrotriazine class called the “glimins” (3). Three Phase III clinical trials were recently completed and strong efficacy was seen in multiple trials (3–5). Imeglimin’s mode of action involves dual effects; to ameliorate insulin resistance and potentiate GSIS (6,7).

Imeglimin has prominent effects to reverse β-cell dysfunction and amplify GSIS: it ameliorates hyperglycemia in models with pancreatic deficient β-cell mass and function including neonatal streptozotocin (N0STZ) diabetic rats and Goto-Kakizaki (GK) rats and increases insulinogenic index during glucose tolerance tests (6); *in vivo* GSIS is enhanced in both lean and high-fat fed rats (8); increased GSIS was seen in hyperglycemic clamps in non-diabetic and N0STZ-diabetic rats (6). In addition, a strictly glucose-dependent effect to enhance insulin secretion was seen with non-diabetic isolated rat islets (8). Moreover, 7 day administration of Imeglimin to Type 2 diabetes patients substantially amplified net GSIS as assessed by hyperglycemic clamp (9).

Given major effects on GSIS, we tested the hypothesis that Imeglimin could acutely and directly impact β-cell dysfunction using islets isolated from Type 2 diabetes animal models (GK and N0STZ-diabetic rats). As an emerging therapeutic option for patients, it is also important to elucidate the mechanism of action. Thus, we conducted a series of studies using islets isolated from GK rats to define effects on pathways leading to GSIS amplification. GK rats are a non-obese Type 2 diabetes model of “isolated” β-cell dysfunction; many features resemble human disease including a loss of first phase insulin secretion, reduced β-cell mass, reduced islet insulin content, inflammation in islets, and impaired islet mitochondrial function (10). Here, we determined that the mechanism of action of Imeglimin was distinct vs. common antidiabetic therapies (metformin or sulphonylureas) and independent from mechanisms mediating the effects of other agents known to affect GSIS (GLP1 receptor agonists or phospholipase C pathway modulators). In contrast, Imeglimin increases NAD^+^ levels in GK rat islets, potentially via the “salvage pathway” involving NAMPT and also increases cellular ATP content, suggesting an improvement in mitochondrial function. Further, we provide evidence suggesting a link, via CD38 and the generation of key NAD^+^ metabolites, between the increased NAD^+^ pool and enhanced intracellular Ca2+ mobilization. These findings implicate a novel mode of action for Imeglimin that could be further leveraged to support the selection of appropriate patients and enhance its clinical utility or to develop improved agents in this new therapeutic class.

## METHODS

### Animals, Islet Isolation, Insulin Secretion and Intracellular Ca2+

Animal studies were conducted at Metabrain Research (Maisons-Alfort, France) according to European guidelines (ETS 123) and were approved by the Ethics Committee. Rats were housed 4 per cage in controlled room (22°C; 12 hour light-dark cycle) with ad libitum access to water and normal chow (A113; Scientific Animal Food and Engineering, AUGY-France). N0STZ rats were obtained by intravenous injection of streptozotocin (100 mg/kg) of rat pups (Charles River) as described (11); 11-12 week-old rats with hyperglycemia and defective GSIS were used (12). Male Wistar rats (11-14 week-old; Charles River) and male GK rats (14 week old; Metabrain Research) were also used.

Rats were anesthetized with i.p. sodium pentobarbital and sacrificed by decapitation. Islets were prepared by injection of collagenase (Sigma) into the pancreatic duct and surgical removal of the pancreas. The pancreas was digested for 9-11 min at 37°C, filtered and rinsed (Hank’s buffer solution containing BSA), and purified with a Ficoll gradient (Sigma) followed by several washes. For static incubations, islets were distributed into 24 well plates; 9-16 wells per group with 6-12 islets per well, depending on the experiment. Islets were incubated for 20-30 min in Krebs Ringer Buffer (KRB) 0.2% BSA with and without test compounds in low (2.8 mM) or high (16.7 mM) glucose (DMSO 0.1% for all conditions) followed by removal of supernatant samples (stored at −20°C until insulin was measured. Selected test agents included Imeglimin (Poxel SA), GLP1, metformin, an imidazoline (13) phospholipase C (PLC) pathway activator (BL11282, Metabrain Research) and a PLC inhibitor (U73122, SIGMA U6756).

For perifusions, islets were distributed (12 per well; 4 well-plates) in KRB containing 5.5 mM glucose and BSA (5 mg/ml) and maintained at 37°C under 95% O_2_/5% CO_2_. In selected studies, islets were loaded with Fura-2-AM (7.5 μM) added to buffer for 1 hour followed by three buffer exchanges. Batches of 8 islets each were placed in a chamber and perifused at 1 ml/min with Hepes-BSA (1mg/ml) buffer alternately containing glucose 2.8mM or 16.7mM with or without test compounds. Perifusate was collected every minute. For intracellular Ca2+, the chamber was placed on the stage of a NIKON TE300 microscope (37°C); individual islets were imaged via excitation at 340nm and 380nm and fluorescence detection (510nm) with a photomultiplier (Photon Technologies International, Princeton, NJ). Intracellular Ca2+ results were expressed as ratio of F340nm/F380nm. Insulin levels were measured via Elisa (Alpco 80-INSRTU-E01 or 80-INSRT-E01).

### Measurement of Intracellular Analytes

For cAMP, GK islets were incubated 30 min in 2.8 mM glucose and then incubated 15 min in 2.8 or 16.7 mM glucose with or without test compounds plus a phosphodiesterase inhibitor (IBMX 1 mM) to prevent cAMP degradation. Supernatants were removed by centrifugation and islets were maintained at −80°C in lysis buffer (Amersham RPN225). cAMP levels were subsequently measured using the same kit.

Dinucleotide content was determined with 20 islets/well in 96 well filter plates; islets were placed in KRB with 16.7mM glucose with or without Imeglimin or nicotinamide (Sigma). Gallotannin was also used where noted (Santa Cruz, K2613). After 20 min, supernatants were removed by centrifugation and islets were stored at −80°C followed by lysis in PBS-dodecyltrimethylammonium bromide solution; NAD^+^ and NADH were determined using Promega kit G9071; NADP^+^ and NADPH were determined using Promega kit G9081.

For ATP and ADP, islets (50 per dish) were stabilized in 5 ml of KRB, 0.2% BSA with glucose 2.8mM for 30 min followed by distribution into 24 well plates (20 islets/well) in KRB 0.2% BSA with glucose 16.7 mM with or without test compounds. After 10 min, islets were transferred to 96 well filter plates and then maintained at −80°C. After lysis (ATP kit buffer), ATP content was measured by luminescence (ATP lite, Perkin Elmer, 6016643); ADP content was measured with a fluorimetric assay (Sigma Aldrich, ref. MAK033).

### NAMPT Activity and Gene Expression

Islets were lysed in 50mM Tris-HCl pH 7.5/0.02% BSA, 0.1% Triton X-100; iNAMPT activity was determined in pools of 60 islets with a colorimetric Cyclex assay kit (Clinisciences, ref. CY-1251). Human recombinant *(E. Coli)* NAMPT activity was measured using the same kit after 60 min. incubation.

Frozen (−80°C) islets (pools of 20) were homogenized followed by extraction and purification (RNAzol kit). RT-PCR measurements employed the AMV reverse transcriptase system (Applied Biosystems 4368814) and Q-PCR reactions (7900HT Fast Real-Time PCR, Applied Biosystems) using primers corresponding to two different exons. Levels of NAMPT mRNA were expressed as increases or decreases in cycle time [Ct] numbers compared to control after normalization to HPRT or β-actin housekeeping genes.

### CD38 Knockdown in Islets

Islets were cultured 24 hours in RPMI medium (11 mM glucose plus inactivated serum, antibiotics, glutamine, 10 mM HEPES) and then placed in 10 cm^2^ plates (100 islets, each), washed in PBS and incubated 15 min on ice in permeabilization buffer (Lyovec 40μl/100 islets/5ml medium, Invitrogen) with siRNA from Origen (10 nM scrambled sequence or 10 nM directed against CD38). Islets were then cultured for 48h before further testing; 15 to 20 wells per group (10 islets/well). Static incubation in 16.7 mM glucose with or without test compounds was followed by removal of supernatant samples for insulin measurements and transfer of islets tubes for RNA extraction as above; CD38 mRNA levels were measured as described above for NAMPT.

### Modulation of cADPR and NAADP Signaling

Islets were distributed (50 per dish) in 5 mL RPMI medium (11 mM glucose), and cultured at 37°C in 95% O_2_ and 5% CO_2_ for 72 hr. For the last 17 hr., high concentration (200μM) Ryanodine (EnzoLife Sciences – Ref. ALX-630-062-M005), was added to selected dishes. After transfer to fresh dishes and incubation for 30 min (KRB/BSA buffer containing 2.8 mM glucose with or without Ryanodine), islets were distributed (6 per well) in 24-well plates in KRB containing 16.7 mM glucose with and without the indicated stimuli or inhibitors that also included cADPR (1 mM; Biolog–Ref. C005-025), NAADP (50 nM; SIGMA N5655), or combinations of two agents. After 20 min. incubation, samples of supernatants were removed and stored at −20°C.

### Statistics

Statistical analyses were performed using a Kruskall-Wallis non parametric one way ANOVA test followed by the Dunn’s post test (GraphPad PRISM4). Where noted, comparison between two conditions was performed using an unpaired Student t-test.

## RESULTS

### Imeglimin Amplifies GSIS in Diseased Rat Islets

β-cell function (GSIS) was impaired (−65% p<0.001) in N0STZ rat islets vs. Wistar control islets (Fig. 1A). GLP1 induced a non-significant trend (+42%) towards increased GSIS in N0STZ islets (Fig. 1B). In low glucose, Imeglimin did not modify insulin secretion; in 16.7mM glucose, increased insulin secretion was observed.

**Fig 1.**
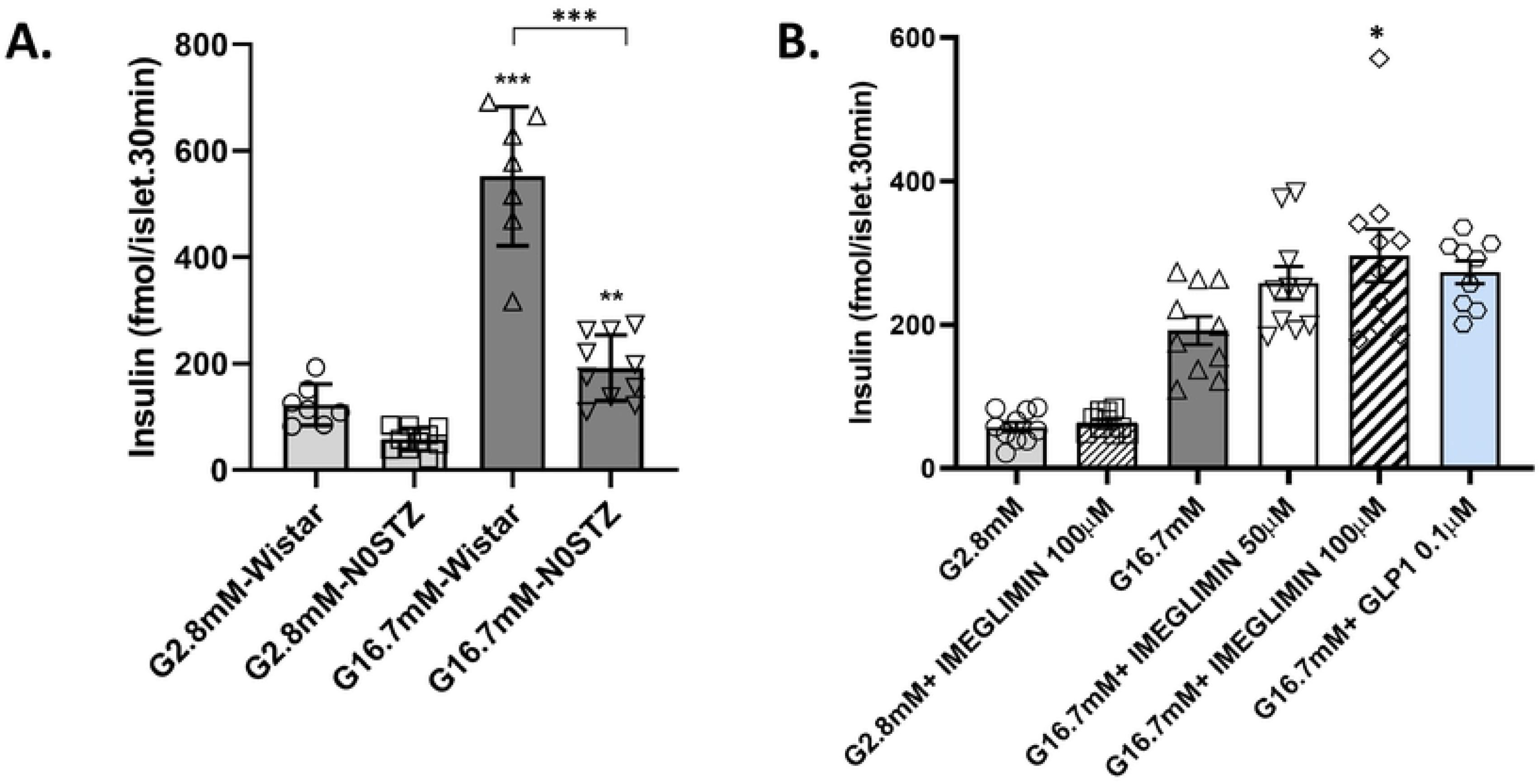
Imeglimin Amplifies Insulin Secretion in Islets from N0STZ Rats. N0STZ Rat Islets vs. Wistar Rat Islets (A). Islets from N0STZ or healthy Wistar rats were incubated in the presence of 2.8mM or 16.7mM glucose. Insulin levels were measured in supernatants after 30 min of incubation. **p<0.01, ***p<0.001 vs. respective low glucose values; mean ± SEM; n=6 wells with 6 islets per well. Effect of Imeglimin and GLP1 on Insulin Secretion from N0STZ Rat Islets (B). Islets from N0STZ rats were incubated in the presence of 2.8m or 16.7mM glucose with or without the tested concentrations of Imeglimin or GLP1 10^-7^M. Insulin levels were measured in supernatants after 30 min of incubation. The effect of Imeglimin at 100 μM was significant, *p<0.05, vs. high glucose alone; mean ± SEM; n=9-10 wells with 6 islets per well (note that when using an unpaired Student t-test, GLP1 also achieved statistical significance, p=0.0054).

GSIS in GK rat islets was markedly impaired vs. a 2-fold response to high glucose in control Wistar islets (Fig. 2A). Imeglimin potentiated GSIS; similar to the results obtained using N0STZ rat islets, Imeglimin was without any effect at low glucose (Fig. S1). A dose-related effect was also evident with a magnitude similar to GLP1 (Fig. 2B). Under the same experimental conditions, we confirmed that metformin could not enhance GSIS (Fig 2C). The effect of 100 μM Imeglimin to ampifly insulin secretion in the presence of high glucose was replicated in 6 additional experiments (S1 Table). Using a perifusion system (Fig. 2D), Imeglimin was also shown to augment GSIS. In this context, the response to high glucose in control GK rat islets was negligible whereas islets from healthy Wistar rats were robustly responsive (Fig. S2). Imeglimin resulted in a partial restoration of GSIS relative to the response noted in Wistar rat islets (compare Fig. 2D and Fig. S2).

**Fig 2.**
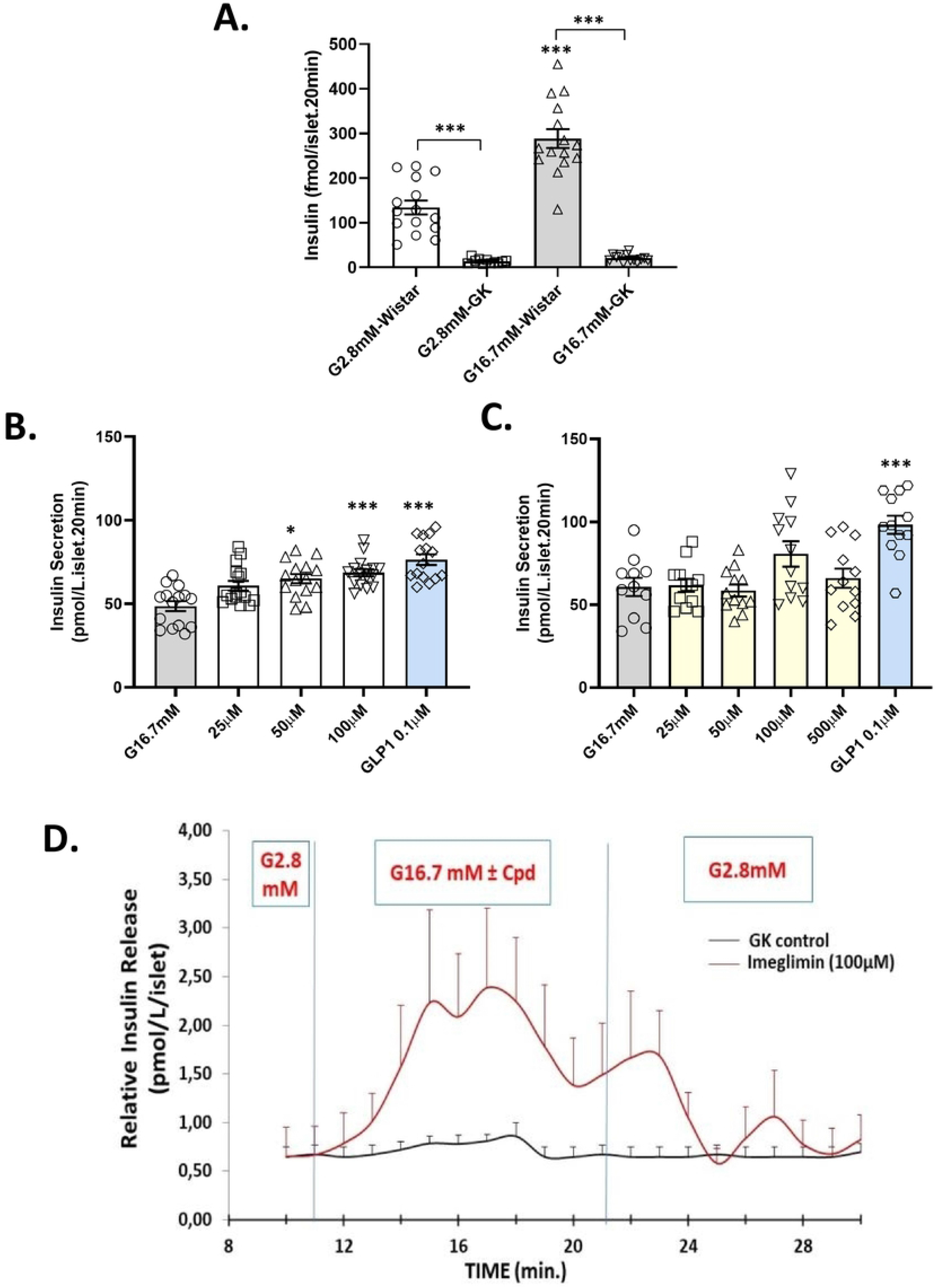
Imeglimin Amplifies Insulin Secretion in Islets from GK Rats. GK Rat Islets vs. (Control) Wistar Rat Islets (A). Islets from GK and Wistar rats were incubated in the presence of glucose 2.8 mM or 16.7 mM. Insulin levels were measured after 20 min of incubation. *p<0.05, **p<0.01, ***p<0.001 vs. respective control value; mean ± SEM; n=6 wells with 6-10 islets per well. Imeglimin (but not Metformin) Amplifies Insulin Secretion from GK Rat Islets: Islets from GK rats were incubated in the presence of high (16.7 mM) glucose (grey bars) or with high glucose plus the indicated concentrations of Imeglimin (B; open bars), metformin (C; yellow bars), or GLP1 as a control (blue bars; panels B and C). Significant increases in mean (± SEM) glucose-stimulated insulin release are noted vs. respective control values; *p<0.05, **p<0.01, ***p<0.001; n=15 to 16 observations per group. Effects of Imeglimin on Kinetics of Insulin Secretion from GK Rat Islets (D). Islets from GK rats were alternately perifused with 2.8 mM glucose for 10 minutes and 16.7 mM glucose with (red curve) or without (black curve) Imeglimin (100μM) for 10 minutes (10 to 20 min) followed by perifusion with 2.8mM for an additional 10 minutes. The insulin levels in the perifusate was measured every minute from 0 min to 30 min. Mean ± SEM insulin levels are shown (data are derived from 4 independent experiments for each group at each time point).

In cadaveric islets derived from a single patient donor with Type 2 diabetes, we also observed an effect (+129%, p<0.05; n=8-10) of Imeglimin (100 μM) to amplify GSIS (Fig. S3).

### Imeglimin’s Actions are Distinct vs. Other Glucose-Dependent Mechanisms

The combination of Imeglimin with GLP1 resulted in trends towards greater GSIS (Fig. S4). These results suggest that Imeglimin and GLP1 may be acting via independent pathways to amplify insulin release. To confirm this hypothesis, we excluded an effect of Imeglimin on cAMP, the classical mediator of GLP1 action, under the same conditions where GLP1 exerted a strong effect (Fig. 3). In β-cells, phospholipase C (PLC) also mediates the potentiation of insulin secretion in response to molecules that include GPR40 (free fatty acid receptor 1) agonists that potentiate GSIS (14). We excluded a role for PLC via use of a specific PLC inhibitor (15)(Fig. S5). These results suggest that Imeglimin and GPR40 agonists act via independent pathways to amplify insulin release.

**Fig 3.**
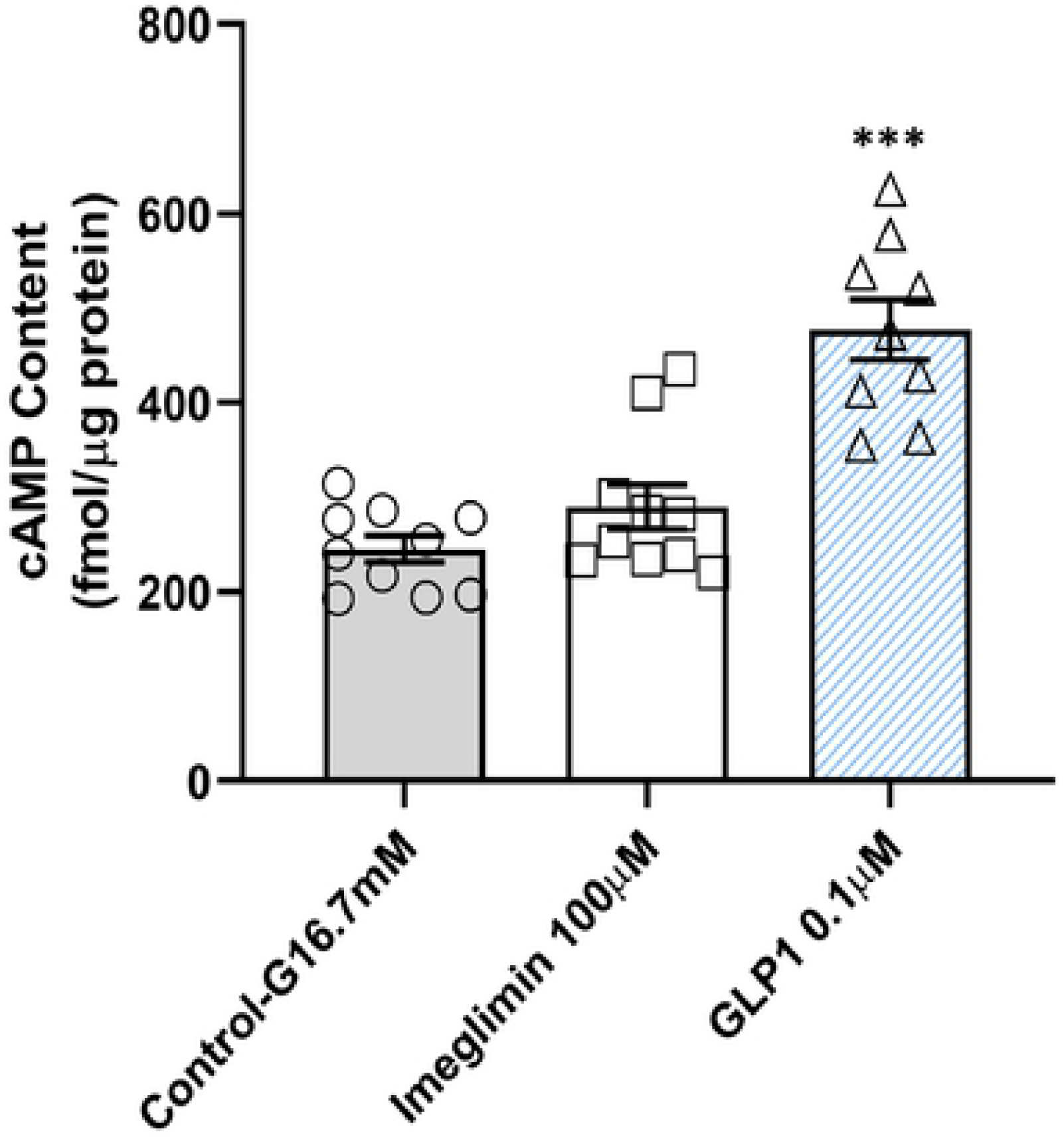
Imeglimin Does Not Increase cAMP Generation in Isolated GK Rat Islets. In the presence of high glucose and the phosphodiesterase inhibitor IBMX, GLP1 (0.1μM) treatment increased the cAMP content of GK islets (+95%, ***p<0.001; n=9). However, Imeglimin (100 μM), produced no effect to increase cAMP under the same conditions. Mean ± SEM values are shown (n=10). An additional independent experiment was also performed; levels of cAMP in each tested condition were not different between the two experiments.

### Imeglimin Modulates Adenine Dinucleotide and ATP Levels

Adenine dinucleotides are known to modulate insulin secretion; we found that both Imeglimin and exogenous nicotinamide induced increases in islet NAD^+^ content and the NAD/NADH ratio under high glucose conditions (Table I). No differences in adenine dinucleotides content were noted when low vs. high glucose alone were compared (data not shown).

**TABLE I.**
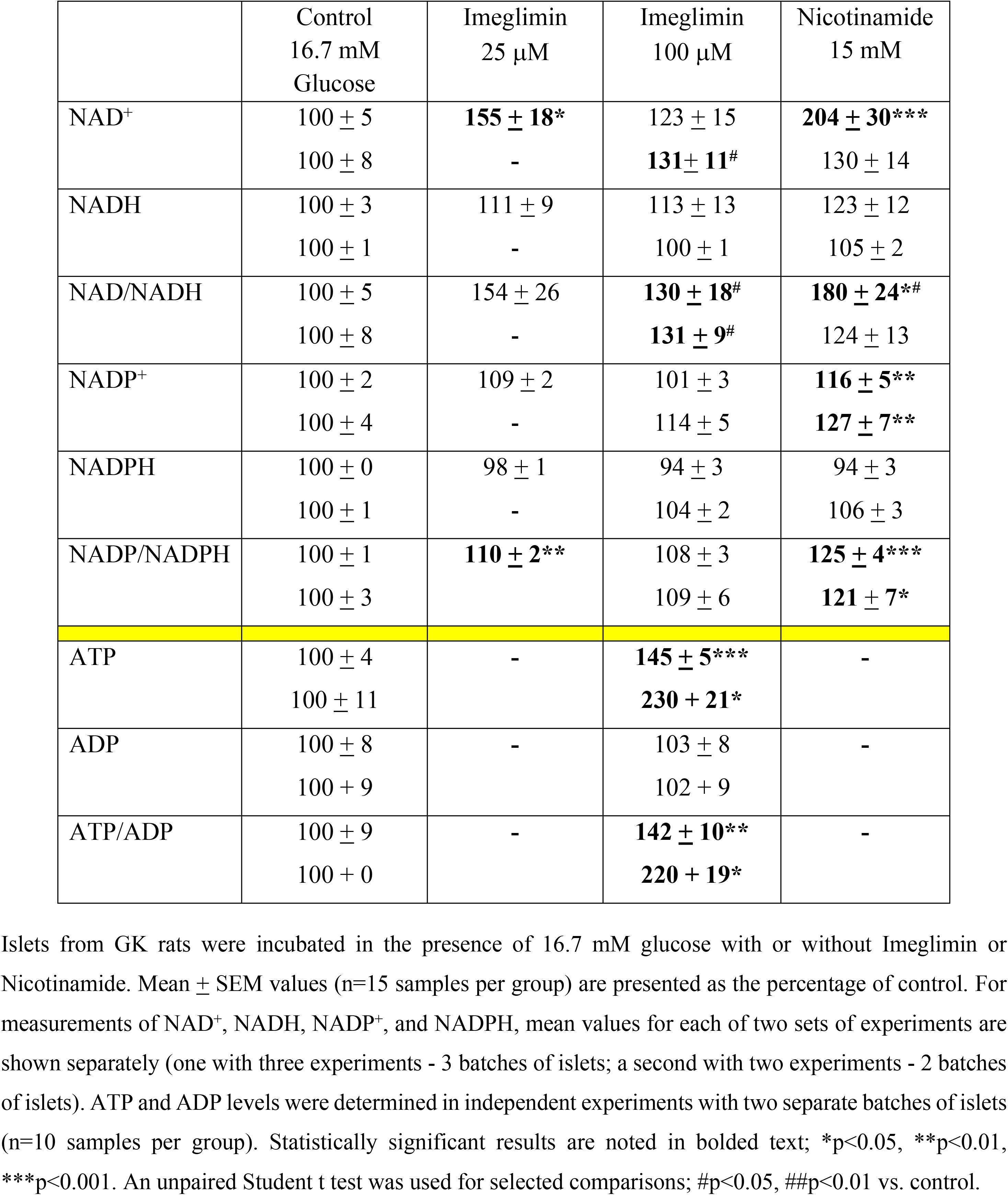
Imeglimin and Nicotinamide Effects on Adenine Dinucleotide and ATP, ADP Content of GK Rat Islets

As NAD^+^ is an essential co-factor for mitochondrial function (16), we also measured ATP levels. The measurement of islet ATP content was validated by assessing the acute (10 min.) effect of exposure to high (16.7 mM) vs. low (2.8 mM) glucose alone; a +47 + 10% increase in ATP was measurable in this context (p<0.05; n=14-16 observations in each group). In the presence of high glucose, Imeglimin significantly increased mean ATP content and the ATP/ADP ratio (Table I). The effect of metformin was also characterized; no such effect was detected with metformin (Fig. S6). To confirm that increases in islet NAD^+^ are sufficient to amplify GSIS in diseased islets, we showed that insulin secretion and NAD^+^ content were increased by exogenous nicotinamide (Fig. S7).

### Increased NAD^+^ via the Salvage Pathway - Increases in NAMPT Expression and Activity

To assess if increases in the NAD^+^ pool are due to enhanced synthesis, we used Gallotannin, an inhibitor of nicotinamide mononucleotide adenylyl transferase (NMNAT), a key enzyme in the NAD^+^ synthetic pathway (17, 18). Gallotannin (10μM) alone had no effect on NAD^+^. As expected, Imeglimin or 15 mM nicotinamide increased NAD^+^ levels (Fig. 4A). With Gallotannin coadministration, NAD^+^ content in Imeglimin treated islets was no longer above control levels and NAD^+^ content in nicotinamide treated islets was partially suppressed. These results suggest that the effect of Imeglimin on NAD^+^ content is mediated by increased synthesis.

**Fig 4.**
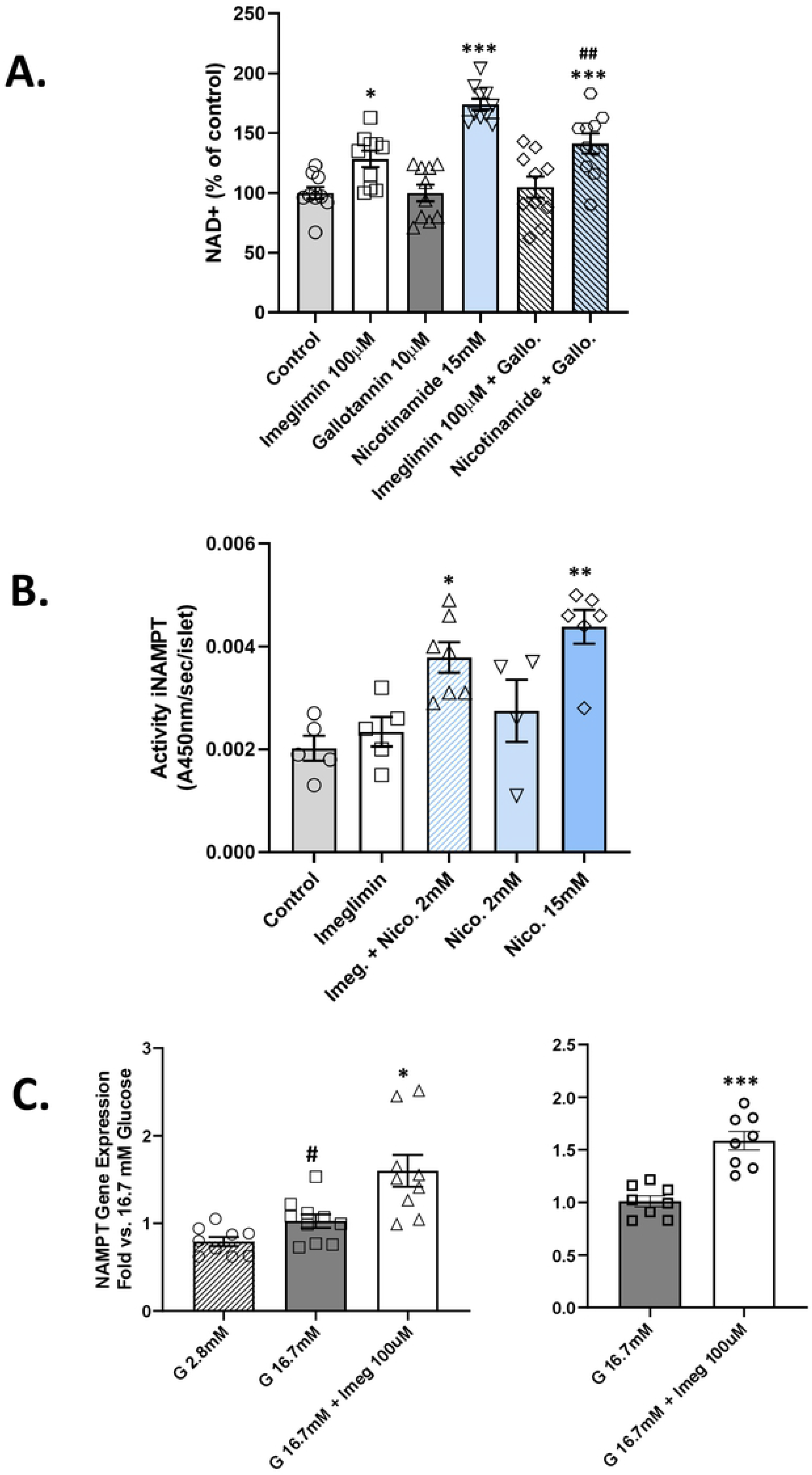
Imeglimin Increases the NAD^+^ Pool Through Increased Synthesis. Gallotannin Effect on NAD^+^ (A). Islets from GK rats were incubated in the presence of 16.7 mM glucose with or without Imeglimin (100μM), or nicotinamide (15mM); compounds were administered alone or in combination with gallotannin (10μM). NAD^+^ was measured after 20 min incubation; mean (n=10 in each group) ± SEM values are shown; *p<0.05, ***p<0.001 vs. Control; ## p<0.01 vs. nicotinamide alone. iNAMPT Activity (B). Islets from GK rats were incubated in the presence of 16.7 mM glucose with or without Imeglimin (100μM), or nicotinamide (2 mM or 15 mM), or the combination of Imeglimin and 2 mM nicotinamide. Intracellular (i) NAMPT activity was then measured; mean ± SEM (n=5-6 per group) values are shown. *p<0.05, **p<0.01 vs. Control. NAMPT mRNA Levels (C). Results from two separate experiments (Right and Left panels) are shown. NAMPT gene expression was determined by RT-PCR in islets from GK rats that were incubated for 30 min. in the presence of 2.8 mM glucose (hatched bar), 16.7 mM glucose (solid bars) or 16.7 mM glucose plus Imeglimin (100μM; open bars). Mean (± SEM; n=9-10 observations per group) levels of NAMPT mRNA are shown as fold vs. 16.7 mM glucose alone; #p<0.05 vs. 2.8 mM glucose; *p<0.05; ***p<0.001 vs. 16.7 mM glucose.

At a low concentration (2 mM), the NAMPT substrate – nicotinamide – appeared to potentiate the effect of Imeglimin on GSIS (+89% vs. +33% with Imeglimin alone; data not shown). Given this result, the activity of intracellular NAMPT, a key enzyme in the NAD^+^ salvage synthesis pathway, was assessed (Fig. 4B). As expected, iNAMPT activity was greater with 15 mM nicotinamide (+117%, p<0.01) and not significantly increased at 2 mM. In the absence of NAMPT substrate (nicotinamide), Imeglimin did not significantly modify iNAMPT activity; however, with 2 mM nicotinamide, iNAMPT activity was increased by Imeglimin (+88%, p<0.05 vs. control). These findings were replicated in an experiment where iNAMPT activity was induced by the combination of Imeglimin (100 μM) and 1 mM nicotinamide (+42%; p<0.05 vs. both control and nicotinamide alone; data not shown). Thus, in the presence of low concentrations of added substrate, Imeglimin leads to increased NAMPT activity. The possible effect of Imeglimin to directly modulate human recombinant NAMPT activity was also assessed. Recombinant NAMPT enzyme activity was not altered by Imeglimin at six different concentrations (Fig. S8).

Since glucose rapidly induces NAMPT expression in isolated human islets (19); the potential for Imeglimin to upregulate NAMPT mRNA was interrogated. High glucose alone modestly induced NAMPT mRNA levels; added exposure to Imeglimin further increased NAMPT mRNA (Fig 4C).

### Imeglimin’s Effects are Distinct vs. Sulphonylureas

Diazoxide opens K^+^-ATP channels to inhibit GSIS (20, 21); sulphonylureas including tolbutamide mediate channel closure and glucose-independent insulin secretion (22). As expected, tolbutamide (and glibenclamide) increased insulin secretion (Fig. 5A; Fig. S9); diazoxide was also shown to inhibit the effect of tolbutamide (Fig 5A). Control experiments also showed that GK rat islets retain the ability to respond to KCl (Fig. S10). Imeglimin’s effect to augment GSIS was unaffected by diazoxide (Fig. 5B). Taken together with the absence of an Imeglimin effect on insulin secretion in low glucose, these results further suggest that Imeglimin’s mode of action is distinct from sulphonylureas and may involve a pathway(s) that is independent of K^+^-ATP channels.

**Fig 5.**
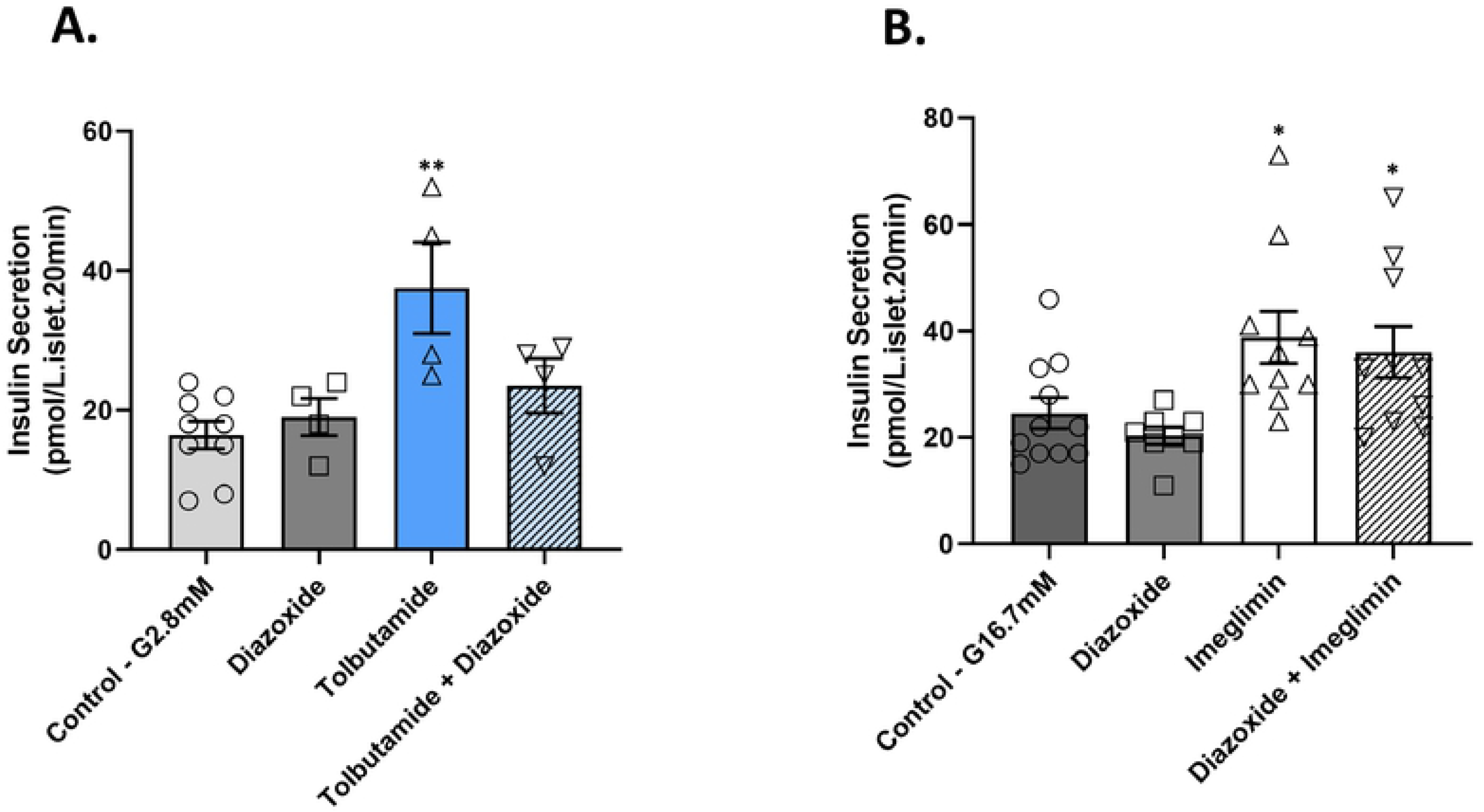
Imeglimin Effect on Insulin Secretion is Resistant to Diazoxide. (A) Islets from GK rats were incubated in low (2.8 mM) glucose with or without diazoxide (400 μM), tolbutamide (500 μM), or a combination of both diazoxide and tolbutamide. (B) GK rat islets were incubated in high (16.7 mM) glucose with or without diazoxide (400 μM), Imeglimin (100 μM), or a combination of both diazoxide and Imeglimin. Samples were obtained after 20 min. and subsequently assayed to determine insulin concentrations; *p<0.05, **p<0.01, vs. respective control value. Mean ± SEM values are shown.

### Potential Role of a CD38-cADPR-Ryanodine Receptor Pathway in NAD^+^ Mediated Mobilization of Intracellular Ca2+

As expected, we also observed that Imeglimin could induce increases in intracellular Ca2+ in response to glucose in GK islets (Fig. 6A). This effect on intracellular Ca2+ was also not observed in low glucose conditions (Fig. S11). We have also observed that glucose-induced Ca2+ mobilization in GK rat islets is impaired by more than 85% vs. Wistar rat islets studied in parallel in a perifusion assay (Fig. S10).

**Fig 6.**
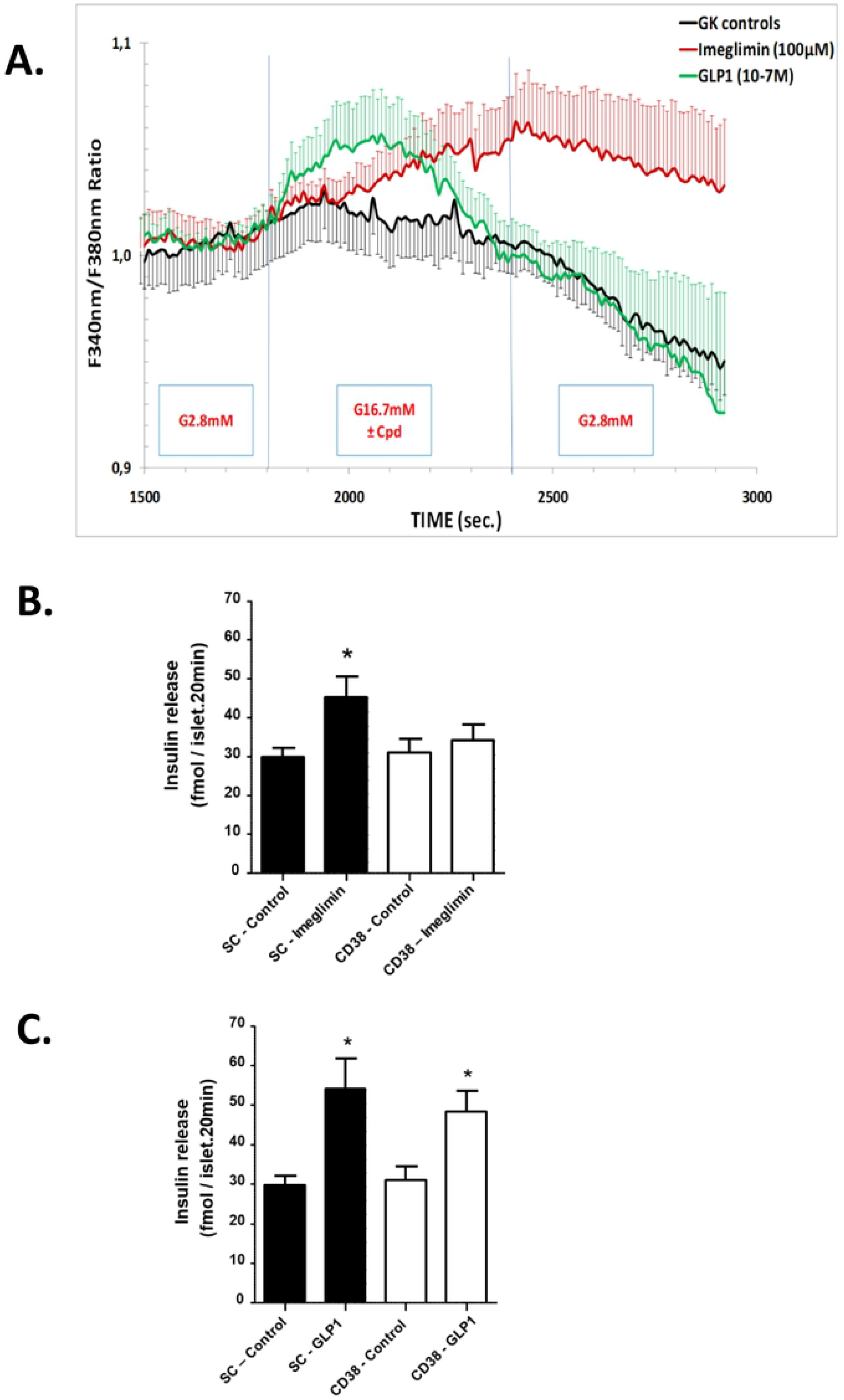
Potential Role of CD38 and NAD^+^ Metabolites to Enhance Insulin Secretion via Increasing Intracellular Ca2+ in Response to Glucose. Measurement of Intracellular Ca2+ in Perifused GK Rat Islets (A). Islets from GK rats were perifused alternately with glucose 2.8 mM and 16.7 mM glucose without treatment for Controls (black curve), with Imeglimin 100 μM (red curve) or with GLP1 0.1 μM (green curve) followed by a third period of perifusion with 2.8 mM glucose alone. Intracellular Ca2+ levels were measured from individual islets by successive excitation at 340 nm and 380 nm and detection of fluorescence emitted at 510 nm every 10 seconds. Results for each of the three groups (control, Imeglimin, GLP1) are derived from 8 experiments with a total of 8 to 10 rats per group (8 rats for control and GLP1 groups, 10 for the Imeglimin group). Insulin Secretion Response to Imeglimin and GLP1 With and Without CD38 Knockdown: Scrambled sequence siRNA control (SC-Control, solid bars) and CD38 siRNA (open bars) transfected GK rat islets were incubated for 20 min. in high (16.7 mM) glucose with or without 100 μM Imeglimin (B) or 0.1 μM GLP (C). Mean ± SEM (n=15-20 per group) insulin release values are shown; *p<0.05 vs. respective control.

NAD^+^ is metabolized to cyclic ADP-ribose (cADPR) and nicotinic acid dinucleotide phosphate (NAADP) via CD38 (cyclic ADP ribose hydrolase). Both metabolites are implicated in mobilizing internal Ca2+ stores, through activation of ER ryanodine receptors in the case of cADPR.

To assess the role of CD38, siRNA-mediated knockdown was employed. CD38 siRNA produced significant and reproducible decreases in CD38 mRNA (from −40% to −49%, p<0.01-0.05; n=14-18 observations per group) vs. control siRNA (data not shown). When CD38 mRNA expression was only moderately reduced, Imeglimin’s effect on GSIS was abolished (Fig. 6B). In contrast, effects of GLP1 treatment were unaffected and there was no effect with scrambled (control) siRNA (Fig. 6C). These results suggest that CD38 is required for the effect of Imeglimin to potentiate GSIS.

Finally, we studied the effects of modulating signaling via cADPR or NAADP on insulin release (Table II). GLP1 and Imeglimin produced expected GSIS effects and exogenous cADPR (1.0 μM) also increased GSIS. cADPR’s effects to enhance Ca2+ mobilization (and GSIS) are reportedly mediated by ryanodine receptors (RyR)(23); thus, high concentration ryanodine was used as a RyR inhibitor. In the presence of 200 μM ryanodine, the effects of either cADPR or Imeglimin to augment GSIS appeared to be abrogated (Table II). However, baseline glucose-stimulated insulin release was also modestly lower in the presence of 200 μM ryanodine vs. without ryanodine, thus complicating the interpretation of these data. Overall, these data suggest a role for cADPR in contributing to Imeglimin’s effects to amplify glucose-stimulated Ca2+ mobilization and insulin secretion.

**TABLE II.** Effects of Modulating cADPR on Glucose-Stimulated Insulin Secretion

| Treatment Group | Insulin Secretion |  | p value(s) |
| --- | --- | --- | --- |
|  | pmol/L.islet.20 min | % of 16.7 mM Glucose Control |  |
| Control 16.7 mM Glucose | 5.1 ± 0.8 | 100 ± 15 | - |
| GLP1 (0.1 µM) | **14.1 ± 1.5** | **274 ± 29** | **<0.001\*** |
| Imeglimin (100 µM) | **7.8 ± 0.7** | **152 ± 13** | **<0.05\*** |
| cADPR (1.0 µM) | **7.3 ± 0.5** | **143 ± 10** | **<0.05\*** |
| Ryanodine (200 µM) | 3.2 ± 0.3 | 63 ± 6 | NS |
| cADPR + Ryanodine | **4.3 ± 0.4** | **84 ± 8** | NS*; **<0.001#** |
| Imeglimin + Ryanodine | **5.4 ± 0.8** | **105 ± 15** | NS*; **<0.05#** |

Islets from GK rats were incubated in the presence of 16.7 mM glucose for 20 min. with or without the indicated compounds as shown; effects pathway inhibition (excess ryanodine) - with or without cADPR or Imeglimin stimulation - are depicted in the lower portion of the table. Mean ± SEM values for insulin released (pmol/L.islet.20 min; also presented as % of 16.7 mM glucose control) are shown (n=8-13 observations per group). Bolded values are statistically significant; p values vs. 16.7 mM glucose control (\*) or vs. the respective single compound in combination treatments (#, first agent listed in Column one) are noted.

## DISCUSSION

The prominant role of β-cell dysfunction in Type 2 diabetes is well established (24–28). Here, we elucidated a novel mechanism by which Imeglimin, a new potential anti-diabetic medication, improves β-cell function – an effect that has been clearly demonstrated *in vivo* in both animal models (6,8) and humans (9).

Imeglimin ameliorates hyperglycemia in rodent models characterized by a primary β-cell defect – STZ-diabetic and GK rats (6). Here, we determined that Imeglimin could acutely and directly enhance GSIS (without any effect in low glucose conditions) with isolated islets from these models. Concentrations where Imeglimin was effective (25-100 μM) are also aligned with human exposure levels (estimated ≈ 50 μM, unpublished; Poxel SA).

Several observations indicate that Imeglimin’s mechanism is distinct vs. other therapeutic approaches. It is important to distinguish the effects of Imeglimin from metformin since in liver there is an apparent overlap with respect to inhibition of gluconeogenesis and the potential to partially inhibit mitochondrial Complex I (6,7). We confirmed that metformin fails to directly potentiate GSIS, consistent with the literature (29,30); in addition, metformin had no effect on GK islet ATP (vs. significant increases with Imeglimin). GLP1 binding to its cognate G-protein coupled receptor induces rapid activation of plasma membrane associated adenylyl cyclase leading to clear increases in cAMP (2,31); Imeglimin had no such effect. Sulphonylureas such as tolbutamide, are secretagogues in both low- and high-glucose; in contrast, the effects of Imeglimin (like GLP1) are only glucose-dependent. We also found that, unlike sulphonylureas, Imeglimin’s effect on GSIS was retained in the presence of diazoxide, a classical β-cell K^+^-ATP channel opener (32). Together with the observed lack of effect on insulin secretion under low glucose conditions in this and prior (8) studies, these findings are consistent with the likelihood of a K^+^-ATP independent mechanism for Imeglimin. Importantly, the GSIS enhancing effects of incretins like GLP1 also involve a diazoxide-resistant K^+^-ATP independent pathway (33). GPR40 agonists and molecules in the imidazoline class have been pursued as GSIS enhancing therapies; these agents operate through PLC activation (14,34) which was also excluded a requirement for Imeglimin’s action.

Mitochondrial dysfunction is a key feature of β-cell dysfunction (35–37); decreases in ATP generation have been described in islets from GK rats and patients with Type 2 diabetes (35, 38–40). We previously showed that Imeglimin can modulate mitochondrial function in liver (7). In islets from healthy rats, Imeglimin was shown to amplify insulin secretion in response to obligate mitochondrial fuels (8). Here, we showed that Imeglimin increased islet ATP levels, an effect that may be consistent with the potential to enhance mitochondrial metabolism. The lack of diazoxide inhibition of Imeglimin’s effect is still compatible with enhanced mitochondrial function since it is well known that additional anaplerotic mitochondrial metabolic cycles also mediate GSIS without requiring downstream K^+^-ATP channel closure (41).

Given its known roles in mitochondrial function, we measured NAD^+^ and demonstrated an increase with Imeglimin, and with nicotinamide, a substrate for NAD^+^ production. Importantly, exogenous nicotinamide was previously shown to enhance GSIS in rodent and human islets (42–44). We confirmed this effect and showed that providing additional substrate for NAD^+^ synthesis - low nicotinamide concentrations – appeared to act in concert with Imeglimin to augment GSIS. These results suggest that pathways emanating from NAD^+^ remain competent in GK islets and may be involved in mediating Imeglimin’s efficacy. NAD^+^ biogenesis occurs via *de novo* synthesis from tryptophan or the salvage pathway from nicotinamide via NAMPT (16,45). Gallotannin, which inhibits NAD^+^ synthesis via both pathways (17,18), was used to provide further results suggesting that Imeglimin’s effect to increase the NAD^+^ pool involves new synthesis of NAD^+^. We also excluded a direct effect of Imeglimin on NAMPT activity *in vitro.* The effect of Imeglimin to induce NAMPT gene expression and activity is intriguing but it is uncertain if this fully accounts for the net increase in NAD^+^ given the short time frame within which these effects were seen. Relevance of the potential role of NAMPT is underscored by studies showing NAMPT expression in β-cells (including human) and that NAMPT haplodeficiency impairs GSIS in mice (19,46).

Increased intracellular Ca2+ is critical for insulin granule exocytosis; Ca2+ sources include both extracellular (via voltage-gated channels in response to K^+^-ATP closure) and intracellular pools (31,48,49). Having observed that Imeglimin can augment Ca2+ mobilization, we assessed a potential link to NAD^+^ generation. In addition to other roles (45,50), metabolism of NAD^+^ by CD38 generates key second messengers – cADPR and NAADP - that are implicated in Ca2+ signaling (45,51). Increases in cADPR, in turn, can activate ryanodine receptors resulting in mobilization of Ca2+ stores from ER (23,49) and this pathway is reportedly operative in pancreatic β-cells (51,52). Our results suggest that Imeglimin’s mechanism is dependent on components of this pathway. However, the efficiency of CD38 knockdown was limited and additional studies will be required to confirm and extend these findings. Although CD38 is described as an ectoenzyme (17), it also exists in an inward orientation and can consume intracellular NAD^+^ (17,53). This pathway is highlighted by increases in islet cADPR and GSIS resulting from β-cell-specific CD38 overexpression in mice (54). However, we acknowledge cADPR’s role in islet function is controversial; especially given an inability to consistently show that cADPR drives Ca2+ release (55). Some of these discrepancies may have resulted from differences in species and methodologies (51).

In assessing the potential role of an NAD^+^ mediated effect to enhance Ca2+ mobilization, our experiments were limited by an inability to measure levels of cADPR in islets from this diabetic rat model, not further interrogating the possible role of NAADP or showing a direct correlation between changes in Ca2+ and the apparent effects of modulating CD38 or RyR. Our studies were also restricted to short time points and may have missed additional, later, effects. There is also a clear need to more precisely define a direct molecular target(s) for Imeglimin including mechanism(s) that may be responsible for induction of NAMPT gene expression.

In summary, we have demonstrated a direct and acute effect of Imeglimin to augment GSIS in diseased islets from a rat model that closely resembles human Type 2 diabetes. The potential mode of action we propose (Fig. 7) involves a pathway leading to increased NAD^+^ content which has been implicated in the regulation of intracellular Ca2+ and is distinct from that employed with other classes of antidiabetic medications including incretin mimetics, sulphonylureas, and biguanides. Additional studies will be required to assess the extent to which pathways implicated in the present studies are also modulated by Imeglimin in human islets. The results reported here are also consistent with existing clinical data where Imeglimin has been shown to enhance GSIS and effectively treat hyperglycemia without any additional risk of hypoglycemia.

**Fig 7.**
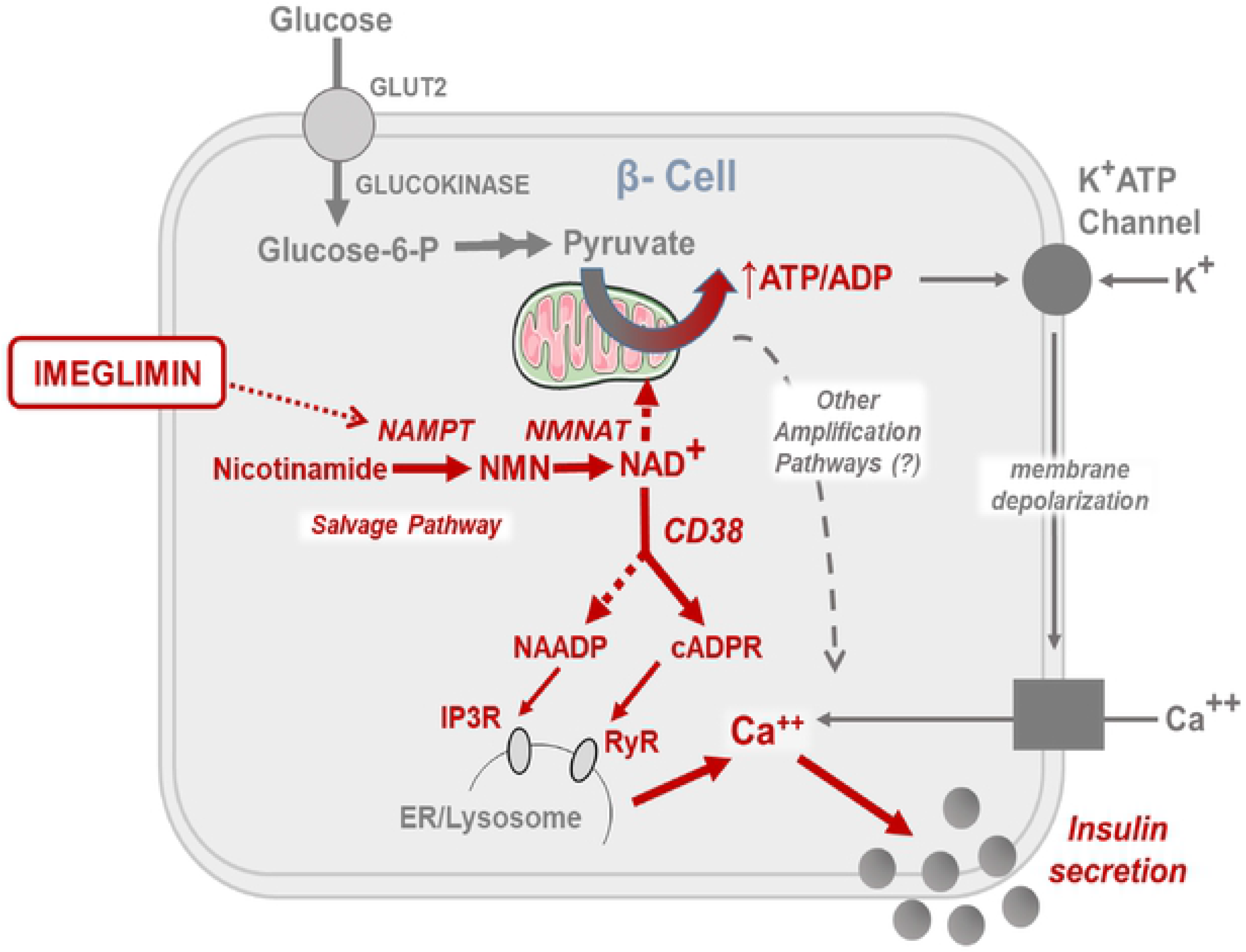
Proposed Model for Mechanism of Imeglimin Action in Islet β-Cells. The effects of Imeglimin in the context of glucose stimulation are highlighted in red (text and arrows).

## ACKNOWLEDGEMENTS

The authors thank Christopher Newgard (Duke University Molecular Physiology Institute, Durham NC) and Johan Auwerx (Laboratory of Integrative Systems Physiology, EPFL, Lausanne, Switzerland) for their helpful comments.

## SUPPORTING INFORMATION CAPTIONS

**Fig. S1 Imeglimin Effects on Insulin Release from GK Rat Islets in Low vs. High Glucose Conditions**

**S1 Table Summary of Additional Experiments Demonstrating Increased GSIS with Imeglimin in GK Rat Islets**

**Fig. S2 Comparison of GSIS in Isolated Islets from Healthy Wistar vs. Diabetic GK Rats**

**Fig. S3 Effect of Imeglimin on GSIS in Isolated Islets from a Patient Donor with Type 2 Diabetes**

**Fig. S4 Effects of Imeglimin on GSIS in GK Rat Islets when Added to Maximal GLP1**

**Fig. S5 Inhibition of Phospholipase C Signaling**

**Fig. S6 Metformin Does Not Affect ATP Levels in Isolated GK Rat Islets**

**Fig. S7 Increases in NAD^+^ Content of GK Rat Islets are Sufficient to Augment Insulin Release**

**Fig. S8 Imeglimin Does Not Modulate the Activity of Recombinant NAMPT**

**Fig. S9 Control Experiments with Diazoxide, Sulphonylureas, KCl**

**Fig. S10 Comparison of Intracellular Ca2+ Responses to Glucose in Wistar vs. GK Rat Islets**

**Fig. S11 Lack of Effect of Imeglimin on Intracellular Ca2+ in the Presence of Low Glucose**

## REFERENCES

1. Kahn SE, Cooper ME, Del Prato S. Pathophysiology and treatment of type 2 diabetes: Perspectives on the past, present, and future. The Lancet 2014; 383:1068–1083.

2. Drucker DJ. Mechanisms of action and therapeutic application of glucagon-like peptide-1. Cell Metab 2018; 27:740–756.

3. Pirags V, Lebovitz H, Fouqueray P. Imeglimin, a novel glimin oral antidiabetic, exhibits a good efficacy and safety profile in type 2 diabetic patients. Diabetes Obes Metab 2012; 14:852–858.

4. Fouqueray P, Pirags V, Inzucchi SE, Bailey CJ, Schernthaner G, Diamant M, et al. The efficacy and safety of Imeglimin as add-on therapy in patients with type 2 diabetes inadequately controlled with metformin monotherapy. Diabetes Care 2013; 36:565–568.

5. Fouqueray P, Pirags V, Diamant M, Schernthaner G, Lebovitz HE, Inzucchi SE, et al. The efficacy and safety of Imeglimin as add-on therapy in patients with type 2 diabetes inadequately controlled with sitagliptin monotherapy. Diabetes Care. 2014; 37:1924–1930.

6. Fouqueray P, Leverve X, Fontaine E, Baquié M, Wollheim C., et al. Imeglimin - A new oral anti-diabetic that targets the three key defects of type 2 diabetes. J Diabetes Metab 2011; 02:126 [doi:10.4172/2155-6156.1000126].

7. Vial G, Chauvin MA, Bendridi N, Durand A, Meugnier E, Madec AM, et al. Imeglimin normalizes glucose tolerance and insulin sensitivity and improves mitochondrial function in liver of a high-fat, high-sucrose diet mice model. Diabetes 2015; 64:2254–2264.

8. Perry RJ, Cardone RL, Petersen MC, Zhang D, Fouqueray P, Hallakou-Bozec S, et al. Imeglimin lowers glucose primarily by amplifying glucose-stimulated insulin secretion in high-fat-fed rodents. Am J Physiol - Endocrinol Metab 2016; 311:E461–E470.

9. Pacini G, Mari A, Fouqueray P, Bolze S, Roden M. Imeglimin increases glucose-dependent insulin secretion and improves β-cell function in patients with type 2 diabetes. Diabetes Obes Metab 2015; 17:541–555.

10. Portha B, Giroix M-H, Tourrel-Cuzin C, Le-Stunff H, Movassat J. The GK rat: a prototype for the study of non-overweight Type 2 diabetes. In: Animal Models in Diabetes Research. Humana Press; 2012. p. 125–159.

11. Portha B, Levacher C, Picon L, Rosselin G. Diabetogenic effect of streptozotocin in the rat during the perinatal period. Diabetes 1974; 23:889–895.

12. Portha B, Picon L, Rosselin G. Chemical diabetes in the adult rat as the spontaneous evolution of neonatal diabetes. Diabetologia 1979; 17:371–377.

13. Efanov AM, Zaitsev S V., Mest HJ, Raap A, Appelskog IB, Larsson O, et al. The novel imidazoline compound BL11282 potentiates glucose-induced insulin secretion in pancreatic β-cells in the absence of modulation of KATP channel activity. Diabetes 2001; 50:797–802.

14. Yamada H, Yoshida M, Ito K, Dezaki K, Yada T, Ishikawa SE, et al. Potentiation of glucose-stimulated insulin secretion by the GPR40-PLC-TRPC pathway in pancreatic β-cells. Sci Rep 2016; 6:2–10.

15. Bleasdale JE, Thakur NR, Gremban RS, Bundy GL, Fitzpatrick FA, Smith RJ, et al. Selective inhibition of receptor-coupled phospholipase C-dependent processes in human platelets and polymorphonuclear neutrophils. J Pharmacol Exp Ther 1990; 255:756–768.

16. Katsyuba E, Mottis A, Zietak M, et al. De novo NAD+ synthesis enhances mitochondrial function and improves health. Nature 2018; 563:354–359.

17. Liu L, Su X, Quinn WJ, Hui S, Krukenberg K, Frederick DW, Redpath P, et al. Quantitative analysis of NAD synthesis-breakdown fluxes. Cell Metab 2018; 27:1067–1080.

18. Petrelli R, Felczak K, Cappellacci L. NMN/NaMN adenylyltransferase (NMNAT) and NAD kinase (NADK) inhibitors: Chemistry and potential therapeutic applications. Curr Med Chem 2011; 18:1973–1992.

19. Kover K, Tong PY, Watkins D, Clements M, Stehno-Bittel L, Novikova L, et al. Expression and regulation of Nampt in human islets. PLoS One. 2013; 8:1–11.

20. Burr IM, Marliss EB, Stauffacher W, Renold AE. Diazoxide effects on biphasic insulin release: “adrenergic” suppression and enhancement in the perifused rat pancreas. J Clin Invest. 1971; 50:1444–1450.

21. MacDonald PE, De Marinis YZ, Ramracheya R, Salehi A, Ma X, Johnson PRV, et al. A KATP channel-dependent pathway within α cells regulates glucagon release from both rodent and human islets of langerhans. PLoS Biol. 2007; 5:1236–1247.

22. Inagaki N, Gonoi T, Clement IV JP, Namba N, Inazawa J, Gonzalez G, et al. Reconstitution of IKATP: An inward rectifier subunit plus the sulfonylurea receptor. Science 1995; 270:1166–1170.

23. Mészáros LG, Bak J, Chu A. Cyclic ADP-ribose as an endogenous regulator of the non-skeletal type ryanodine receptor Ca2+ channel. Nature 1993; 364:76–79.

24. Kahn SE, Zraika S, Utzschneider KM, Hull RL. The beta cell lesion in type 2 diabetes: There has to be a primary functional abnormality. Diabetologia 2009; 52:1003–1012.

25. Deng S, Vatamaniuk M, Huang X, Doliba N, Lian MM, Frank A, et al. Structural and functional abnormalities in the islets isolated from Type 2 diabetic subjects. Diabetes 2004; 53:624–632.

26. Del Guerra S, Lupi R, Marselli L, Masini M, Bugliani M, Sbrana S, et al. Functional and molecular defects of pancreatic islets in human type 2 diabetes. Diabetes. 2005; 54:727–735.

27. Butler AE, Janson J, Bonner-Weir S, Ritzel R, Rizza RA, Butler PC. β-cell deficit and increased β-cell apoptosis in humans with type 2 diabetes. Diabetes 2003; 52:102–110.

28. Rahier J, Guiot Y, Goebbels RM, Sempoux C, Henquin JC. Pancreatic β-cell mass in European subjects with type 2 diabetes. Diabetes Obes Metab 2008; 10 (SUPPL. 4):32–42.

29. DeFronzo RA, Barzilai N, Simonson DC. Mechanism of metformin action in obese and lean non-insulin dependent diabetic subjects. J Clin Endocrinol Metab 1991; 73:1294–1301.

30. Rena G, Hardie DG, Pearson ER. The mechanisms of action of metformin. Diabetologia. 2017:1577–1585.

31. Meloni AR, Deyoung MB, Lowe C, Parkes DG. β-cells: Mechanism and glucose dependence. Diabetes Obes Metab 2013; 15:15–27.

32. Proks P, Reimann F, Green N, Gribble F, Ashcroft FA. Sulfonylurea stimulation of insulin secretion Diabetes 2002; 51:S368–S376.

33. Yajima H, Komatsu M, Schermerhorn T, Aizawa T, Kaneko T, Nagai M, et al. cAMP enhances insulin secretion by an action on the ATP-sensitive K+ channel-independent pathway of glucose signaling in rat pancreatic islets. Diabetes. 1999; 48:1006–1012.

34. Efendic S, Efanov AM, Berggren P, Zaitsev SV. Two generations of insulinotropic imidazoline compounds. Diabetes 2002; 51:S448–S454.

35. Anello M, Lupi R, Spampinato D, Piro S, Masini M, Boggi U, et al. Functional and morphological alterations of mitochondria in pancreatic beta cells from type 2 diabetic patients. Diabetologia 2005; 48:282–289.

36. Ma ZA, Zhao Z, Turk J. Mitochondrial dysfunction and β-cell failure in type 2 diabetes mellitus. Exp Diabetes Res 2012; 2012: Article ID 703538; doi:10.1155/2012/703538.

37. Haythorne E, Rohm M, van de Bunt M, Brereton MF, Tarasov AI, Blacker TS, et al. Diabetes causes marked inhibition of mitochondrial metabolism in pancreatic β-cells. Nat Comm 2019; 10. Available from: http://dx.doi.org/10.1038/s41467-019-10189-x

38. Sasaki M, Fujimoto S, Sato Y, Nishi Y, Mukai E, Yamano G, et al. Reduction of reactive oxygen species ameliorates metabolism-secretion coupling in islets of diabetic GK rats by suppressing lactate overproduction. Diabetes 2013; 62:1996–2003.

39. Giroix MH, Vesco L, Portha B. Functional and metabolic perturbations in isolated pancreatic islets from the GK rat, a genetic model of noninsulin-dependent diabetes. Endocrinology 1993; 132:815–822.

40. Tsuura Y, Ishida H, Okamoto Y, Kato S, Sakamoto K, Horie M, et al. Glucose sensitivity of ATP-sensitive K+ channels is impaired in b-cells of the GK rat: A new genetic model of NIDDM. Diabetes 1993; 42:1446–1453.

41. Jensen MV, Joseph JW, Ronnebaum SM, Burgess SC, Sherry AD, Newgard CB. Metabolic cycling in control of glucose-stimulated insulin secretion. 2008; Am J Physiol Endocrinol Metab 295: E1287–E1297.

42. Deery DJ, Taylor KW. Effects of azaserine and nicotinamide on insulin release and nicotinamide-adenine dinucleotide metabolism in isolated rat islets of Langerhans. Biochem. J. 1973, 134: 557–563

43. Otonkoski T, Beattie GM, Mally MI, Ricordi C, Hayek A. Nicotinamide is a potent inducer of endocrine differentiation in cultured human fetal pancreatic cells. J Clin Invest 1993; 92:1459–1466.

44. Zawalich WS, Dye ES, Matschinsky FM. Nicotinamide modulation of rat pancreatic islet cell responsiveness in vitro. Horm Metab Res 1979;11:469–471.

45. Cantó C, Menzies KJ, Auwerx J. NAD+ metabolism and the control of energy homeostasis: A balancing act between mitochondria and the nucleus. Cell Metab 2015; 22:31–53.

46. Revollo JR, Körner A, Mills KF, Satoh A, Wang T, Garten A, et al. Nampt/PBEF/Visfatin regulates insulin secretion in β Cells as a systemic NAD biosynthetic enzyme. Cell Metab 2007 7:363–375.

47. Graves TK, Hinkle PM. Ca2+-induced Ca2+ release in the pancreatic β-cell: Direct evidence of endoplasmic reticulum Ca2+ release. Endocrinology 2003; 144:3565–3574.

48. Kang G, Holz GG. Amplification of exocytosis by Ca2+-induced Ca2+ release in INS-1 pancreatic β cells. J Physiol 2003; 546:175–189.

49. Lee HC, Walseth TF, Bratt GT, Hayes RN, Clapper DL. Structural determination of a cyclic metabolite of NAD+ with intracellular Ca2+-mobilizing activity. J Biol Chem 1989; 264:1608–1615.

50. Katsyuba E, Romani M, Hofer D, Auwerx J. NAD+ homeostasis in health and disease. Nature Metab 2020; 2:9–31.

51. Zhao Y, Graeff R, Lee HC. Roles of cADPR and NAADP in pancreatic cells. Acta Biochim Biophys Sin (Shanghai). 2012; 44: 719–729.

52. Takasawa S, Nata K, Yonekura H, Okamoto H. Cyclic ADP-ribose in insulin secretion from pancreatic β cells. Science 1993; 259:370–373.

53. Zhao YJ, Zhu WJ, Wang XW, Zhang L-H, Lee HC. Determinants of the membrane orientation of a calcium signaling enzyme CD38. Biochim Biophys Acta 2015; 1853:2095–2103.

54. Kato I, Takasawa S, Akabane A, Tanaka O, Abe H, Takamura T, et al. Regulatory role of CD38 (ADP-ribosyl cyclase/cyclic ADP-ribose hydrolase) in insulin secretion by glucose in pancreatic β cells: Enhanced insulin secretion in CD38-expressing transgenic mice. J Biol Chem. 1995; 270:30045–30050.

55. Islam MS, Berggren PO. Cyclic ADP-ribose and the pancreatic beta cell: Where do we stand? Diabetologia. 1997; 40:1480–1484.

